## Supplemental Methods for "Microbes display broad diversity in cobamide preferences"

*Media and growth conditions*

*Sinorhizobium meliloti* was cultured in M9 medium supplemented with 0.2% sucrose, 10 ng/ml biotin, and 10 μM cobalt chloride (1). *S meliloti* Rm1021 Δ*nrdJ cobD*::*gus* $Gm^R$ *bhbA*::Tn*5* pMSO3-*nrdAB*(*E. coli*) (KM168) and Δ*metH cobD*::*gus* $Gm^R$ *bhbA*::Tn*5* (KM622) were used for MetH- and NrdJ-dependent growth assays, respectively. KM168 was precultured in M9 sucrose medium supplemented with 0.1 mg/ml L-Methionine (Met) and MetH-dependent growth assays were performed in the same medium lacking Met. KM622 was precultured in two steps; first it was grown in M9 sucrose medium supplemented with 1 mg/ml Met and 100 nM cyanocobalamin (CNCbl) for 48 hours, then an aliquot was diluted 100-fold in M9 sucrose supplemented with Met and grown for an additional 24 hours. NrdJ-dependent growth assays were performed in M9 sucrose supplemented with Met. All media were additionally supplemented with 20 μg/ml gentamycin and cultures were grown at 30°C with aeration.

For *Escherichia coli* MetH-dependent growth assays, *E. coli* MG1655 Δ*metE* was used for testing natively expressed *metH*, while *E. coli* MG1655 Δ*metE* Δ*metH* was used when expressing *metH* orthologs from the pETmini plasmid. All strains were precultured in M9 glucose (0.2%) medium supplemented with 0.1 mg/ml Met and growth assays were performed in the same medium lacking Met. Cultures of strains carrying pETmini-*metH*(*E. coli*) and pETmini-*metH*(*V. cholerae*) were supplemented with 25 μg/ml kanamycin. For *E. coli* EAL-dependent growth assays, *E. coli* MG1655 Δ*metH* was precultured in ethanolamine medium (2) supplemented with 0.02% ammonium chloride and EAL-dependent growth assays were performed in the same medium lacking ammonium chloride.

*Bacteroides thetaiotaomicron* VPI 5482 (3) was precultured in BHIS liquid medium (4) with growth assays performed in Varel-Bryant glucose defined medium without Met (4). *Ruminococcus gnavus* ATCC 29149 was precultured in M2 medium from Tramontano *et al*. (5) with growth assays performed in M2 medium lacking Met. *B. thetaiotaomicron* and *R. gnavus* were cultured at 37°C in an anaerobic chamber (Coy Laboratory Products Inc.) under an atmosphere of approximately 10% $CO_2$, 3% $H_2$, and 87% $N_2$.

*Bacillus subtilis* strain KK1401 (*Em his nprE18 aprE3 eglS*Δ102 *bglT*/*bglS*ΔEV *lacA*::P*xylA*-*comK loxP*-Pveg-*btuFCDR queG*::*loxP amyE*::Pveg-*metH*(*Priestia megaterium*)) (6), which expresses *P. megaterium* DSM319 *metH*, was precultured in LB for 3 hours to harvest cells during early exponential phase ($OD_{600}$ approximately 0.25). MetH-dependent growth assays were performed in Spizizen minimal medium supplemented with 0.5% glucose and 0.03% L-Histidine (7). *B. subtilis* was grown at 37°C with aeration.

*Growth Assays*

Precultures of *S. meliloti*, *E. coli*, *B. subtilis*, and *B. thetaiotaomicron* were inoculated with individual colonies following growth on LB agar supplemented with 2.5 mM $MgSO_4$ and 2.5 mM $CaCl_2$ (*S. meliloti*), LB agar (*E. coli* and *B. subtilis*), or solid BHIS (*B. thetaiotaomicron*). Solid media were supplemented with 10 μM CNCbl and antibiotics when necessary. Cells harvested from preculture media were washed 1-3 times with either 0.85% saline or growth assay media. Growth assays with *S. meliloti*, *E. coli*, and *B. subtilis* were performed in 384-well plates (Nunc, 265202) and incubated in a BioTek Synergy 2 (medium shaking speed) or Tecan Spark microplate reader (linear shaking, 1 mm amplitude, 1440 rpm) and $OD_{600}$ readings were recorded periodically. Plates were sealed with Breathe-Easy (Diversified Biotech). Growth assays with *B. thetaiotaomicron* and *R. gnavus* were performed in an anaerobic chamber in 384-well plates. Plates were sealed with non-breathable seals, which were removed prior to $OD_{600}$ readings taken with a Tecan Infinite M1000 Pro microplate reader. The starting $OD_{600}$ values were as follows: *S. meliloti*, 0.02; *E. coli* (MetH), 0.004; *E. coli* (EAL), 0.02; *B. subtilis*, 0.01, *B. thetaiotaomicron*, 0.01; *R. gnavus*, 0.02. $EC_{50}$ values and 95% confidence intervals were calculated using Graphpad Prism (Dose-response – Stimulation; [Agonist] vs. response – Variable slope (four parameters)).

*Strain and plasmid construction*

*S. meliloti* Rm1021 Δ*nrdJ bhbA*::Tn*5 cobD*::*gus* $Gm^R$ pMSO3-*nrdAB*(*E. coli*) (KM168) was constructed via two sequential phage M12 transductions (8) of *bhbA*::Tn*5* (9) and *cobD*::*gus* $Gm^R$ (10) into Rm1021 Δ*nrdJ* pMSO3-*nrdAB*(*E. coli*) (11). *S. meliloti* Rm1021 Δ*metH bhbA*::Tn*5 cobD*::*gus* $Gm^R$ (KM622), was constructed via two sequential phage M12 transductions of *bhbA*::Tn*5* and *cobD*::*gus* $Gm^R$ into Rm1021 Δ*metH.* To construct the Δ*metH* strain, 5′ and 3′ flanking regions of the *metH* gene were cloned into pK18*mobsacB* and integrated into the Rm1021 chromosome (12). Neomycin-resistant exconjugants were grown on solid medium containing 10% sucrose to select for a second crossover to create an unmarked *metH* deletion (12), which was confirmed by PCR.

*E. coli* Δ*metE*::$kan^R$ and Δ*metH*::$kan^R$ mutations from the Keio collection (13) were introduced by P1 transduction into *E. coli* strain MG1655 (14). Kanamycin resistance cassettes were removed by introducing plasmid pCP20 carrying the Flp recombinase as described (15), leaving the FRT site in place of the coding sequences. *metH* from *E. coli* MG1655 and *V. cholerae* O1 biovar El Tor str. N16961 were cloned into pETmini (16) via Gibson assembly.

*Cobamide reagents*

Cobamides were used in their cyano- forms for all experiments. Cyanocobalamin was purchased from MilliporeSigma. All other cyanocobamides were extracted from bacterial cultures, purified, and quantified as previously described (17, 18). Benzimidazolyl and phenolyl cobamides were produced in *Sporomusa ovata*, and purinyl cobamides were produced in *Propionibacterium acidi-propionici*.


1. Maniatis T, Fritsch EF, Sambrook J. 1989. Molecular cloning: a laboratory manual. Cold spring harbor laboratory press.
2. Scarlett FA, Turner JM. 1976. Microbial metabolism of amino alcohols. Ethanolamine catabolism mediated by coenzyme $B_{12}$-dependent ethanolamine ammonia-lyase in *Escherichia coli* and *Klebsiella aerogenes*. J Gen Microbiol 95:173-6. doi:10.1099/00221287-95-1-173.

3. Mishra S, Imlay JA. 2013. An anaerobic bacterium, *Bacteroides thetaiotaomicron*, uses a consortium of enzymes to scavenge hydrogen peroxide. Mol Microbiol 90:1356-71. doi:10.1111/mmi.12438.
4. Liu H, Shiver AL, Price MN, Carlson HK, Trotter VV, Chen Y, Escalante V, Ray J, Hern KE, Petzold CJ, Turnbaugh PJ, Huang KC, Arkin AP, Deutschbauer AM. 2021. Functional genetics of human gut commensal *Bacteroides thetaiotaomicron* reveals metabolic requirements for growth across environments. Cell Rep 34:108789. doi:10.1016/j.celrep.2021.108789.
5. Tramontano M, Andrejev S, Pruteanu M, Klunemann M, Kuhn M, Galardini M, Jouhten P, Zelezniak A, Zeller G, Bork P, Typas A, Patil KR. 2018. Nutritional preferences of human gut bacteria reveal their metabolic idiosyncrasies. Nat Microbiol 3:514-522. doi:10.1038/s41564-018-0123-9.
6. Kennedy KJ, Widner FJ, Sokolovskaya OM, Innocent LV, Procknow RR, Mok KC, Taga ME. 2022. Cobalamin Riboswitches Are Broadly Sensitive to Corrinoid Cofactors to Enable an Efficient Gene Regulatory Strategy. mBio 13:e0112122. doi:10.1128/mbio.01121-22.
7. Anagnostopoulos C, Spizizen J. 1961. Requirements for Transformation in *Bacillus Subtilis*. J Bacteriol 81:741-6. doi:10.1128/jb.81.5.741-746.1961.
8. Finan TM, Hartweig E, LeMieux K, Bergman K, Walker GC, Signer ER. 1984. General transduction in *Rhizobium meliloti*. J Bacteriol 159:120-4. doi:10.1128/jb.159.1.120-124.1984.
9. Charles TC, Cai GQ, Aneja P. 1997. Megaplasmid and chromosomal loci for the PHB degradation pathway in *Rhizobium* (*Sinorhizobium*) *meliloti*. Genetics 146:1211-20. doi:10.1093/genetics/146.4.1211.
10. Campbell GR, Taga ME, Mistry K, Lloret J, Anderson PJ, Roth JR, Walker GC. 2006. *Sinorhizobium meliloti bluB* is necessary for production of 5,6-dimethylbenzimidazole, the lower ligand of $B_{12}$. Proc Natl Acad Sci U S A 103:4634-9. doi:10.1073/pnas.0509384103.
11. Taga ME, Walker GC. 2010. *Sinorhizobium meliloti* Requires a Cobalamin-Dependent Ribonucleotide Reductase for Symbiosis With Its Plant Host. Mol Plant Microbe Interact 23:1643-54. doi:10.1094/MPMI-07-10-0151.
12. Schäfer A, Tauch A, Jäger W, Kalinowski J, Thierbach G, Pühler A. 1994. Small mobilizable multi-purpose cloning vectors derived from the *Escherichia coli* plasmids pK18 and pK19: selection of defined deletions in the chromosome of *Corynebacterium glutamicum*. Gene 145:69-73. doi:10.1016/0378-1119(94)90324-7.
13. Baba T, Ara T, Hasegawa M, Takai Y, Okumura Y, Baba M, Datsenko KA, Tomita M, Wanner BL, Mori H. 2006. Construction of *Escherichia coli* K-12 in-frame, single-gene knockout mutants: the Keio collection. Mol Syst Biol 2:0008. doi:10.1038/msb4100050.
14. Silhavy TJ, Berman ML, Enquist LW. 1984. Experiments with Gene Fusions. Cold Spring Harbor Laboratory Press, Cold Spring Harbor, NY.
15. Datsenko KA, Wanner BL. 2000. One-step inactivation of chromosomal genes in *Escherichia coli* K-12 using PCR products. Proc Natl Acad Sci U S A 97:6640-5. doi:10.1073/pnas.120163297.
16. Mok KC, Sokolovskaya OM, Nicolas AM, Hallberg ZF, Deutschbauer A, Carlson HK, Taga ME. 2020. Identification of a Novel Cobamide Remodeling Enzyme in the Beneficial Human Gut Bacterium *Akkermansia muciniphila*. mBio 11. doi:10.1128/mBio.02507-20.
17. Mok KC, Hallberg ZF, Taga ME. 2022. Purification and detection of vitamin $B_{12}$ analogs. Methods Enzymol 668:61-85. doi:10.1016/bs.mie.2021.11.023.
18. Sokolovskaya OM, Mok KC, Park JD, Tran JLA, Quanstrom KA, Taga ME. 2019. Cofactor Selectivity in Methylmalonyl Coenzyme A Mutase, a Model Cobamide-Dependent Enzyme. mBio 10:e01303-19. doi:10.1128/mBio.01303-19.
