## Supplemental Table 1 for "Microbes display broad diversity in cobamide preferences"

| | Cbl | [Bza]Cba | [5-MeBza]Cba | [5-OMeBza]Cba | [Ade]Cba | [2-MeAde]Cba | [*p*-Cre]Cba | [Phe]Cba |
|---|---|---|---|---|---|---|---|---|
| ***S. meliloti*** **(MetH)** (Δ*nrdJ cobD*::*gus bhbA*::Tn*5* pMSO3-*nrdAB*(*E. coli*)) | 3.481 (2.548 - 4.672) | 14.47 (12.76 - 16.32) | | | 1077 (949 - 1216) | 740.4 (< 808.5) | 2159 (> 1971) | > 5000 |
| ***S. meliloti*** **(NrdJ)** (Δ*metH cobD*::*gus*, *bhbA*::Tn*5*) | 88.69 (68.58 - 116.2) | 223.2 (142.3 - 564.3) | | | > 2500 | 1830 (1223 - 7914) | NG | NG |
| ***E. coli*** **(EAL)** (Δ*metH*) | 3.676 (ND) | 1.069 (0.9621 - 1.182) | 2.176 (> 2.006) | 1.442 (< 1.634) | 1.042 (> 0.937) | 0.946 (> 0.847) | NG | |
| ***E. coli*** **(native MetH)** (Δ*metE*) | 0.047 (0.037 - 0.059) | 0.069 (0.060 - 0.080) | 0.048 (0.040 - 0.056) | 0.058 (0.050 - 0.066) | 0.222 (0.208 - 0.237) | 0.094 (0.085 - 0.105) | 10.87 (> 10.01) | > 25 |
| ***E. coli*** **(*E. coli* MetH plasmid)** (Δ*metE* Δ*metH* pETmini-*metH*(*E. coli*) | 0.042 (0.035 - 0.049) | 0.064 (0.057 - 0.073) | 0.044 (0.037 - 0.052) | 0.052 (0.046 - 0.059) | 0.180 (0.168 - 0.193) | 0.094 (0.086 - 0.102) | 4.816 (4.414 - 5.270) | 5.382 (4.723 - 6.113) |
| ***E. coli*** **(*V. cholerae* MetH plasmid)** (Δ*metE* Δ*metH* pETmini-*metH*(*V. cholerae*) | 0.047 (0.041 - 0.053) | 0.074 (0.063 - 0.085) | 0.053 (0.048 - 0.059) | 0.072 (0.065 - 0.080) | 51.75 (47.11 - 56.92) | 0.171 (0.154 - 0.188) | 7.715 (7.020 - 8.568) | 5.672 (5.206 - 6.177) |
| ***B. subtilis*** **(*P. megaterium* MetH)** (Δ*queG*::Pveg-*btuFCDR* Δ*metE*::*loxP* *amyE*::Pveg-*metH*(*P. megaterium*)) | 0.161 (0.141 - 0.184) | 1.273 (< 6.495) | 0.187 (0.169 - 0.208) | 0.496 (0.441 - 0.557) | NG | NG | 17.49 (15.99 - 19.21) | 19.18 (16.52 - 22.35) |
| ***B. thetaiotaomicron*** **(MetH)** | 0.219 (0.205 - 0.234) | 0.788 (0.755 - 0.823) | 0.049 (0.046 - 0.053) | 0.195 (0.184 - 0.207) | 1.728 (1.604 - 1.861) | 0.030 (0.027 - 0.033) | NG | NG |
| ***R. gnavus*** **(MetH)** | 0.039 (0.032 - 0.048) | 0.056 (0.045 - 0.072) | 0.049 (0.039 - 0.063) | 0.139 (0.112 - 0.186) | 0.121 (0.101 - 0.147) | 0.078 (0.065 - 0.093) | > 2.5 | > 5 |

**Table S1. $EC_{50}$ values calculated from data in Figure 1 B-J.** $EC_{50}$ values (nM) and 95% confidence intervals (in parentheses) are given for each growth condition shown in Figure 1B-J (four-parameter non-linear fit in GraphPad Prism (v9.5.1)). Genotypes are given for engineered strains. NG, no growth; empty, not measured. A greater than (>) value for $EC_{50}$ is an estimate for cultures that failed to reach saturation at any cobamide concentration tested. Greater than (>) and less than (<) symbols in confidence intervals were used when upper or lower bounds could not be determined, respectively; ND represents a confidence interval in which both upper and lower bounds could not be determined.
